# Emergence of mpox in Guinea: sporadic clade IIa cases and a clade IIb upsurge

**DOI:** 10.64898/2026.03.03.709059

**Authors:** Jacob Camara, Nils Peter Petersen, Fara Raymond Koundouno, Giuditta Annibaldis, Kaba Kourouma, Barré Soropogui, Sarah Ryter, Saa Lucien Millimono, Moussa Condé, Emily Victoria Nelson, Carolina van Gelder, Mia Le, Eugène Kolie, Karifa Kourouma, Tamba Elie Millimouno, Fernand M’Bemba Tolno, Faya Moriba Kamano, Mamadou Bhoye Keita, Saa Antoine Bongono, Facely Tolno, Ousmane Traoré, Sory Conde, Alimou Camara, Philippe Lemey, Stephan Günther, Sophie Duraffour, Sanaba Boumbaly

**Affiliations:** Centre de recherche en Virologie - Laboratoire des Fièvres Hémorragiques Virales de Guinée (CRV-LFHVG), Conakry, Guinea; Bernhard Nocht Institute for Tropical Medicine (BNITM), Hamburg, Germany; German Center for Infection Research (DZIF), partner site Hamburg–Lübeck– Borstel–Riems, Hamburg, Germany; Laboratoire des Fièvres Hémorragiques Virales de Guéckédou (LFHV-GKD), Guéckédou, Guinea; Institut National de Santé Publique (INSP), Conakry, Guinea; University of Hamburg, Institute for Computational Systems Biomedicine, Hamburg, Germany; Direction préfectorale de la santé de Macenta, Macenta, Guinea; Centre de traitement épidémiologique (CTEpi) de Nongo, Conakry, Guinea; Agence Nationale de Sécurité Sanitaire (ANSS), Conakry, Guinea; Department of Microbiology, Immunology and Transplantation, Rega Institute, KU Leuven, Leuven, Belgium

**Author notes:** Equal contribution first authors (Jacob Camara, Nils Peter Petersen, Fara Raymond Koundouno, Giuditta Annibaldis). Equal contribution last authors (Sophie Duraffour, Sanaba Boumbaly). Corresponding author: Nils Peter Petersen, Bernhard-Nocht-Institute for Tropical Medicine, Bernhard-Nocht-Str. 74, 20359 Hamburg, Germany.

## Abstract

**Background:** Mpox virus (MPXV) has caused recurrent outbreaks in West Africa. However, Guinea had not previously reported laboratory-confirmed cases or MPXV genomic data.

**Methods:** Suspected cases were identified in the N’Zérékoré region as well as in the Conakry (Sept 2024– Dec 2025) and confirmed by real-time PCR in regional and central laboratories, respectively. Whole-genome sequencing using nanopore technology was performed in-country, followed by phylogenetic and time-scaled evolutionary analyses.

**Findings:** The first mpox case was clinically diagnosed in September 2024 in the N’Zérékoré region and laboratory-confirmed by the prefectural laboratory in Guéckédou. In total, seven cases were confirmed in Forest Guinea, of which five complete or almost complete MPXV genomes were recovered. All belonged to MPXV clade IIa. Genomic divergence, ancestral dating, and low APOBEC3-associated mutation frequencies were consistent with multiple independent zoonotic spillover events. In June 2025, one of the first mpox cases of an unfolding outbreak was confirmed in Conakry. Whole genome sequencing revealed MPXV clade IIb lineage G.1. By December 2025, the number of laboratory-confirmed mpox cases nationwide increased to 2,151. A total of nine outbreak strains were sequenced, all belonging to Clade IIb. The genomes clustered with contemporaneous genomes from Sierra Leone and showed high APOBEC3-associated mutation frequencies, suggesting sustained human-to-human transmission in the region.

**Interpretation:** These data demonstrate simultaneous circulation of MPXV clade IIa and IIb strains in Guinea, likely resulting from zoonotic spillover and human-to-human transmission, respectively. Decentralised diagnostics and in-country sequencing facilitated rapid case confirmation and genomic surveillance, highlighting the importance of these critical capacities for outbreak preparedness and response.

**Funding:** German Federal Ministry of Health and the German Center for Infection Research (DZIF).

## Introduction

Mpox, caused by mpox virus (MPXV), has historically been endemic in parts of Central and West Africa, with transmission typically resulting from zoonotic spillover and limited human-to-human spread.^1,2^ MPXV clade I is persistently circulating in Central Africa, whereas clade II is endemic in West Africa.^1^ Within clade II, both clade IIa and clade IIb have been associated with zoonotic spillover events. However, clade IIb has additionally demonstrated sustained human-to-human transmission, culminating in the rapid global spread of mpox in 2022 and prompting the World Health Organization (WHO) to declare a Public Health Emergency of International Concern (PHEIC) in July 2022.^3^ During this global expansion, clade IIb genomes accumulated APOBEC3-associated mutational signatures, which have been interpreted as genomic markers of sustained human-to-human transmission.^4^ The first PHEIC was lifted in May 2023, but rapid increases in reported cases due to the spread of new clade I lineages led to a second PHEIC declaration in August 2024.^3^ In West Africa, several countries, including Guinea’s neighbors Liberia, Sierra Leone, and Côte d’Ivoire, have documented mpox cases, whereas Guinea had previously not reported laboratory-confirmed infections.^2,5,6^

Here, we report the first confirmed mpox cases in Guinea, detected in 2024 through national diagnostic and genomic surveillance, and present their genomic characterization, marking the emergence of mpox in the country and the subsequent upsurge in cases in 2025.

## Methods

### Ethical approval and Nagoya

This work is approved by the National Ethics Committee of Guinea (CNERS) under the number 009/CNERS/25 enabling the retrospective analysis of anonymized diagnostics surveillance data. This work is also part of the Nagoya permit number 006/2023/PN.

### Samples and diagnostics

Human samples consisted of vesicular fluid, pustular fluid or crusts collected as part of the in-country mpox virus (MPXV) surveillance efforts and sent to a network of reference laboratories for real-time polymerase chain reaction (PCR) processing. The laboratories involved in the report included the *Laboratoire des fièvres Hémorragiques Virales de Guéckédou* (LFHV-GKD) located in Guéckédou, and the *Centre de Recherche en Virologie* (CRV), as well as the *Institute National de Santé Publique*, both based in Conakry. Viral DNA was extracted from patient samples using the DNeasy Blood & Tissue Kit (Qiagen, Germany) according to the manufacturer’s instructions. PCR was performed using both the RealStar® Zoonotic Orthopoxvirus PCR Kit 1.0 and the FlexStar® Monkeypox virus PCR Detection Mix 1.5 (altona Diagnostics, Germany). Note that samples processed in September 2024 were also analyzed following a PCR protocol as in Scaramozzino et al.^7^ until the FlexStar® Monkeypox virus PCR essay was implemented in October 2024.

### Untargeted nanopore sequencing

Since the COVID-19 pandemic, the CRV laboratory is equipped with a sequencing facility and thus PCR-positive MPXV samples were processed with nanopore sequencing. DNA was extracted and PCR-tested as described above and sequenced. The DNA eluates were thus directly used for library preparation using the Native Barcoding Kit 24 V14 (SQK-NBD114.24) from Oxford Nanopore Technologies (ONT, United Kingdom) according to the manufacturer’s instructions. The libraries were sequenced on R10.4.1 flow cells for at least 40 hours on the Mk1C (ONT) device.

### Genome assembly

Sequencing reads were basecalled using Guppy version 6.3.7 with the high accuracy model. Genomes were assembled using a custom in-house pipeline (https://github.com/opr-group-bnitm/mopx). First, reads were mapped to the reference genome using Minimap2 version 2.28,^8^ Structural variants were called using Sniffles2 version 2.6.2,^9^ followed by small variant calling and consensus generation using Medaka version 2.0.1 (ONT), resulting in a first-pass draft genome. For scaffolding, *de novo* assemblies were generated with both Canu version 2.2 and Flye version 2.9.5.^10,11^ Contigs were iteratively placed against the draft genome using Minimap2. End-to-end reconstruction of MPXV genome was made by using Flye contigs to identify turning points in contigs that span both strands as MPXV has a covalently closed double-stranded DNA genome. Subsequent polishing was performed using Medaka to improve base-level accuracy. Terminal hairpin structures of approximately 50 nucleotides in length are imperfectly complementary and were manually inspected and corrected. Genomic positions with sequencing coverage below a threshold were masked with ‘N’. The default threshold was 20x. For some cases (PX782069, PX782070, PX782071, PX782072, PX990728, PX990729), the threshold was lowered to 10x and read in the regions with coverage 10x to 20x were manually reviewed. Finally, the final consensus sequences were reviewed and manually corrected where necessary and deposited under the accession numbers PV993193, PV993194, PX088143, PX088144, PX088145, PX088146, PX990729, PX990730, PX782071, PX782072, PX782069, PX990731, PX782070, PX990728. The reference genomes KJ642616.1 and PV993193 were used as for clade IIa assemblies, and NC_063383.1 and PX088143 for clade IIb.

### Phylogenetic analysis

Nextclade was used for an initial clade assignment to each genome.^12^ Two background datasets were assembled to analyse clade IIa and clade IIb genomes separately by downloading full genomes assigned to the respective clades from Pathoplexus.^13^ Because the clade IIb genomes described in this study clustered within lineage G.1, a sub-lineage of lineage A.2, the clade IIb dataset was restricted to lineage A.2 genomes, including sub-lineages A.2, A.2.1, A.2.2, A.2.3, and G.1. Sequences identified as outliers, characterized by unusually long phylogenetic branches and requiring additional curation, as well as sequences lacking required metadata, were manually removed from both datasets.

Multiple sequence alignment was performed using squirrel.^14^ One outgroup genome sequence was added to each alignment to root the maximum likelihood trees. For clade IIa the clade IIb reference genome NC_063383.1 was used. For clade IIb we chose the first clade IIa sequence detected in Guinea (PV993193). Maximum likelihood phylogenetic analysis was performed using IQ-TREE version 3.0.1 with ModelFinder,^15,16^ ancestral state reconstruction and ultra-fast bootstrapping with UFBoot with 1000 bootstrap iteration during each run.^17^ The models JC, HKY, TN93, and GTR were tested with both the discrete Gamma model (G) and the FreeRate model (E), and with empirical base frequencies (F) as well as estimated base frequencies (FU). The best-fit model according to BIC was used. For clade IIa this was GTR with a four-category Gamma model and empirical base frequencies. For clade IIb this was HKY with a four-category Gamma model and empirical base frequencies. Branches were annotated with APOBEC3-related SNPs (TC to TT and reverse complement GA to AA) based on ancestral state reconstruction.

For clade IIa, bayesian phylogenetic analysis was conducted using BEAST 2 version 2.6.1.^18^ The presence of temporal signal was confirmed using TempEST version 1.5.3, ^19^ and a root-to-tip randomization analysis. A correlation coefficient of 0.82 and a R^2^ value of 0.67 between root to tip divergence and collection data were observed. A randomization with 100,000 random permutations yielded a P value of 0 indicating a clear temporal signal. For the BEAST run, the GTR substitution model, empirical base frequencies, and gamma-distributed rate heterogeneity across four categories were chosen based on the model selection results from IQ-Tree2. A relaxed log-normal molecular clock and a coalescent constant population prior were used. Six independent MCMC runs were performed for 150 million steps each, sampling every 10,000 steps, using random seeds (3, 5, 7, 11, 13, 17). Convergence was verified in Tracer,^20^ with all runs showing ESS > 2000 after discarding 10% burn-in. LogCombiner was used to merge samples from the runs with 10% burn-in removed. TreeAnnotator generated a maximum clade credibility tree using median heights.

To visualize phylogenies and geography baltic version 0.3.0 and cartopy version 0.25.0 were used. ^21,22^

## Results

### Detection and genomic characterization of MPXV clade IIa in Guinea (2024–2025)

The first laboratory-confirmed mpox case in Guinea was identified on Sept 2, 2024, in Macenta prefecture, N’Zérékoré region, in a child. Infection was confirmed by real-time PCR at the *Laboratoire des Fièvres Hémorragiques Virales de Guéckédou* (LFHV-GKD) in Guéckédou and subsequently verified at the *Centre de Recherche en Virologie* (CRV) in Conakry. Whole-genome nanopore sequencing performed at CRV identified the virus as mpox virus (MPXV) clade IIa. Between September 2024 and September 2025 overall seven laboratory-confirmed mpox cases were detected in the prefectures of Macenta and Guéckédou. Genomes were successfully recovered from five of them (**Table S1**).

Phylogenetic analysis showed that all five Guinea genomes belonged to MPXV clade IIa. Together with two recently reported genomes from Sierra Leone (Pathoplexus accessions: PP_0049Q0N.2 and PP_002XKQ3.2),^23^ these sequences formed a distinct sub-clade within clade IIa that was separate from previously described genomes from Liberia, Ghana, and Côte d’Ivoire (**Figure 1A**). The closest available relatives of this sub-clade were historical genomes sampled in Europe and the USA between 1958 and 1968, of which the closest was a sequence obtained during a mpox outbreak at Rotterdam Zoo in 1965 (GenBank accession: KJ642614.1^24,25^).

**Figure 1.**
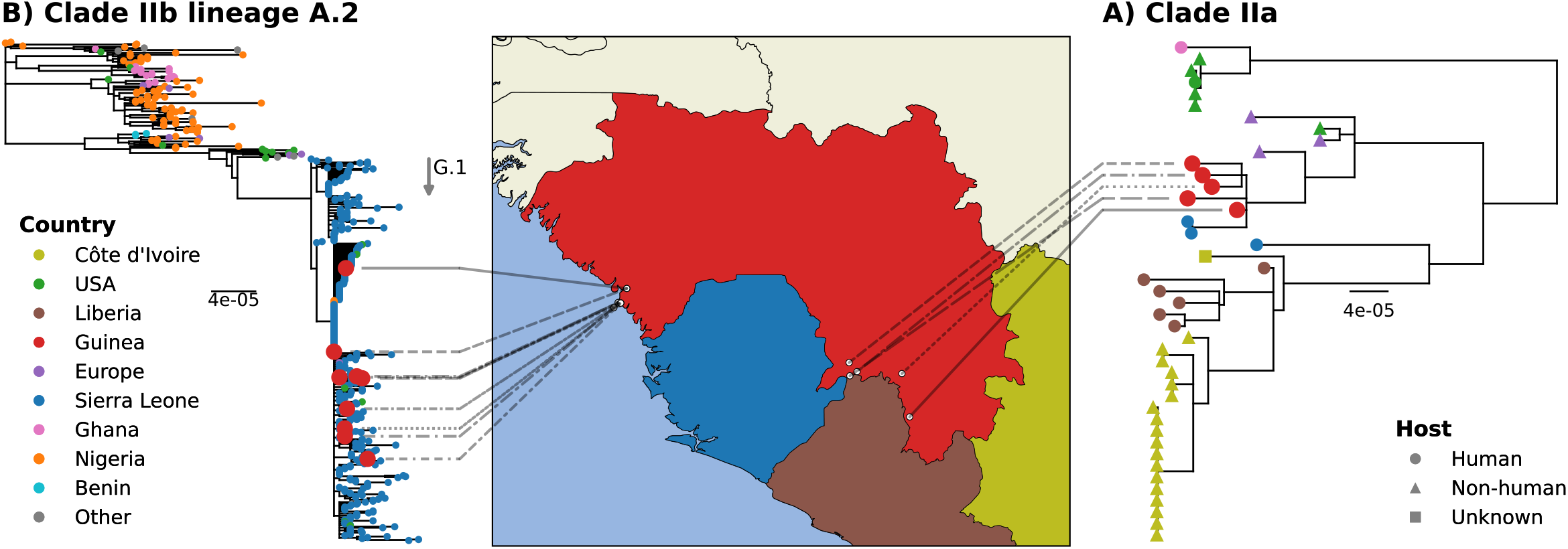
Phylogenetic placement of mpox virus genomes detected in Guinea. Maximum likelihood phylogenies show the position of genomes generated in this study within MPXV clades IIa and IIb. (A) Five genomes obtained from mpox cases detected between 2024 and 2025 in the N’Zérékoré region of Guinea cluster within clade IIa and form part of a distinct sub-clade together with recent genomes from Sierra Leone. U.S. genomes shown at the top are linked to an outbreak associated with rodents imported from Ghana. (B) Genomes generated from cases detected in and around Conakry during the 2025 outbreak cluster with sequences from the contemporaneous outbreak in Sierra Leone and belong to clade IIb lineage G.1, supporting regional transmission across national borders.

Bayesian time-scaled phylogenetic analysis estimated the most recent common ancestor (MRCA) of the clade IIa sub-clade containing Guinea and Sierra Leone genomes to around 2004 (95% highest posterior density [HPD] 2004–2017) (**Figure 2**). The closest related pair of Guinea genomes shared a common ancestor dated to approximately 2013 (95% HPD 2003– 2021), indicating substantial genetic divergence among sampled cases.

**Figure 2.**
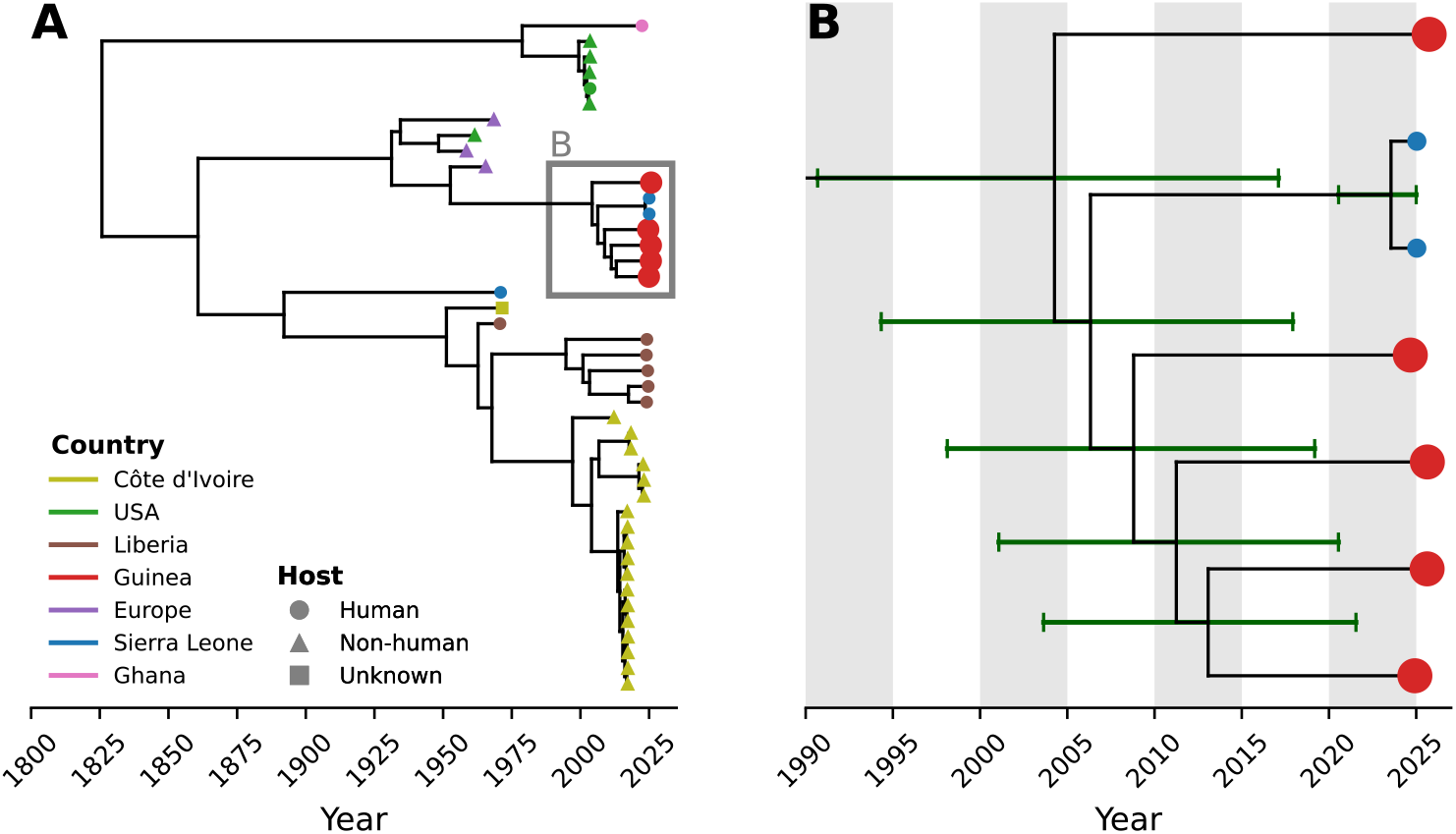
Time-scaled phylogeny of clade IIa mpox virus genomes. (A) Maximum clade credibility trees reconstructed from clade IIa genomes. (B) Expanded view of the sub-clade containing genomes from Guinea and recently reported genomes from Sierra Leone. Node ages and corresponding 95% highest posterior density (HPD) intervals are indicated by green bars. Even the closest Guinea genomes share a common ancestor dating to around 2013 (95% HPD: 2003–2021), consistent with substantial divergence among cases and supporting multiple independent spillover events.

The human APOBEC3 enzyme introduces a characteristic mutational signature in the MPXV genome (TC to TT and GA to AA substitutions). Previous studies have shown that during sustained human transmission, these APOBEC3-associated variants accumulate at an elevated rate of about six mutations per year, compared with a much slower background evolutionary rate of roughly one substitution every three years.^4^ Consequently, these mutations can serve as a signature of sustained human-to-human transmission. However, branch-specific ancestral reconstruction identified only limited APOBEC3-associated mutations among the clade IIa genomes from Guinea. APOBEC3-associated substitutions represented 0/12, 0/2, 0/10, 1/8, and 2/6 of total SNPs across the five clade IIa genomes (**Figure 3**), supporting the view that the corresponding viruses have not yet extensively circulated in the human population, if at all.

**Figure 3.**
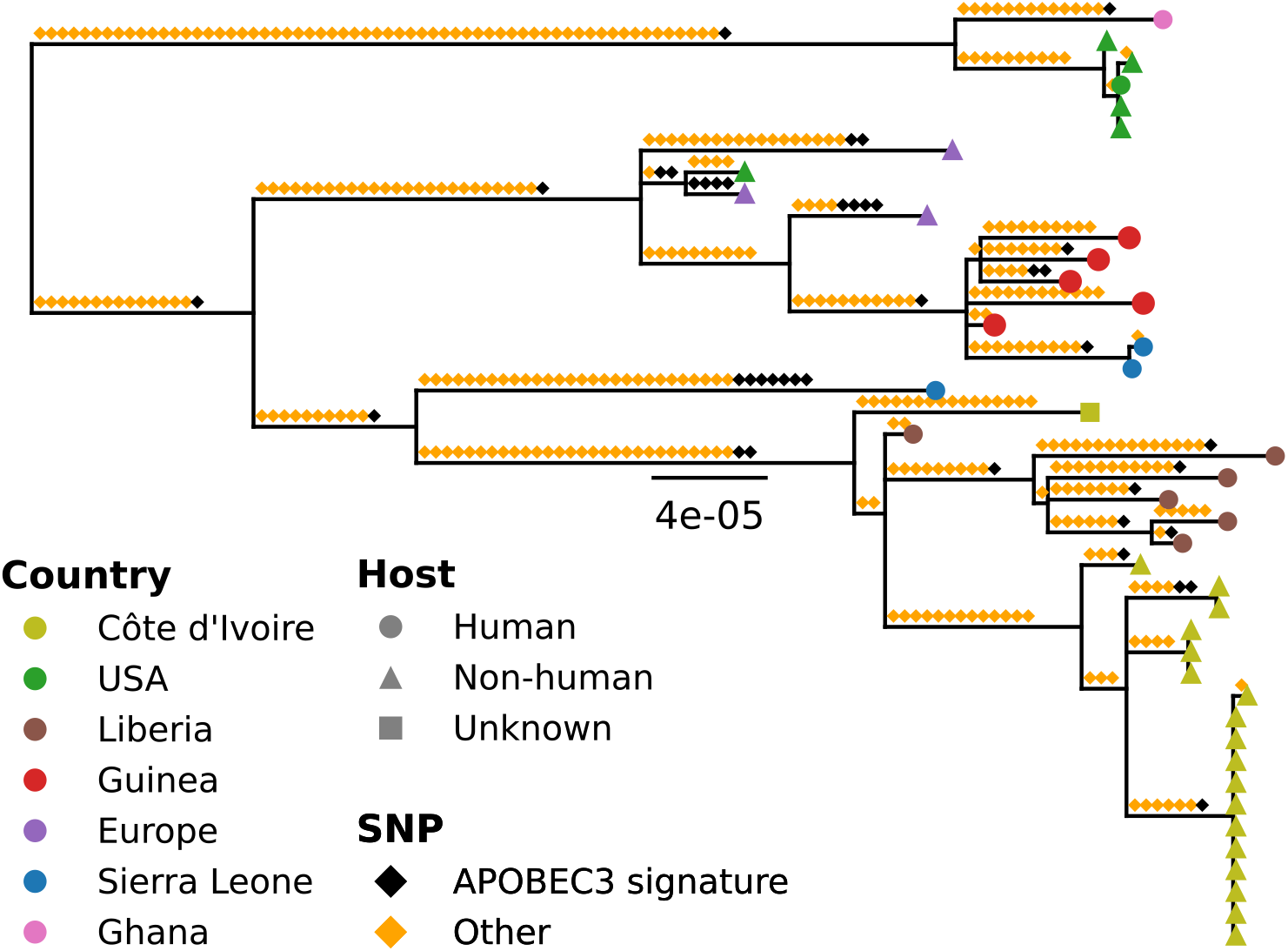
Clade IIa phylogeny annotated with APOBEC3 mutational signatures. Branches are annotated with the number of APOBEC3-associated mutations relative to the total number of SNPs observed. Guinea clade IIa genomes show few APOBEC3-signature mutations (0/12, 0/2, 0/10, 1/8, and 2/6 SNPs), consistent with limited evidence for sustained human-to-human transmission and supporting independent zoonotic introductions.

### Genomic analysis of the 2025 MPXV clade IIb outbreak in Guinea

On 3 June 2025, the National Public Health Institute (INSP) of Guinea detected an mpox case in Kaloum, Conakry, in an adult man, representing the third confirmed mpox case in Guinea at that time. Whole-genome sequencing at CRV assigned the virus to MPXV clade IIb lineage G.1, a sub-lineage of A.2 (**Figure 1B**). By December 31, 2025, a total of 2,151 laboratory-confirmed mpox cases had been reported nationwide in Guinea (including cases from all clades).^26^ Of these 728 were diagnosed at CRV (**Figure S1, Table S2**). In total, nine samples yielded full-length genomes, representing approximately 1·2% of confirmed cases processed at CRV during the study period.

Phylogenetic analysis showed that genomes generated from Guinea during the 2025 outbreak clustered closely with contemporaneous genomes from Sierra Leone within lineage G.1 (**Figure 1B**). The Guinea genomes did not form a distinct monophyletic cluster and bootstrap support was low, and therefore did not provide support for a single cross-border transmission chain between the two countries (**Figure 4**).

**Figure 4.**
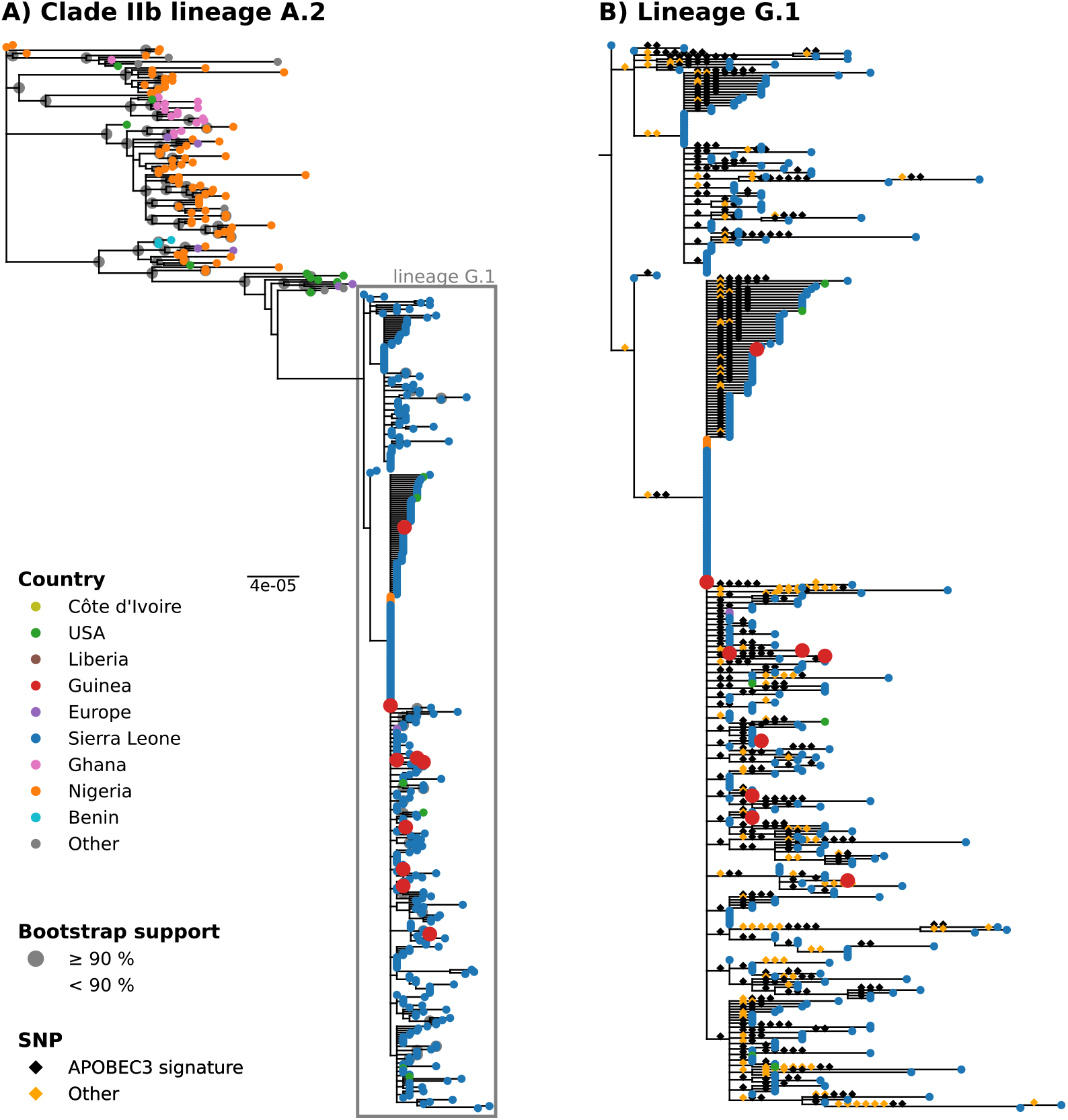
Clade IIb lineage A.2 phylogeny. (A) All clade IIb genomes from Guinea clustered within sub-lineage G.1. (B) Expanded view of the lineage G.1 subtree associated with an outbreak in Sierra Leone. The high frequency of APOBEC3-signature SNPs is consistent with sustained human-to-human transmission.

In contrast to the clade IIa genomes, all clade IIb genomes contained a high frequency of APOBEC3-associated substitutions, suggesting prolonged circulation within the human population prior to sequencing. APOBEC3-signature SNPs accounted for 13/14 substitutions along terminal branches leading to the Guinea genomes and 543/694 substitutions across the entire lineage G.1 subtree (**Figure 4**).

## Discussion

This study describes the first laboratory-confirmed mpox cases in Guinea and demonstrates concurrent circulation of two different MPXV clades. Genomic analyses revealed two distinct molecular epidemiological patterns: genetically diverse clade IIa infections in the N’Zérékoré region presumably resulting from zoonotic spillover and an outbreak of clade IIb lineage G.1 infections consistent with regional human transmission dynamics.

The substantial divergence among clade IIa genomes, together with MRCA estimates predating case detection by more than a decade, suggests that these infections are unlikely to be epidemiologically linked with a common transmission chain. Instead, the observed diversity is consistent with multiple independent zoonotic spillover events. This interpretation is further supported by the low number of APOBEC3-associated mutations, which contrasts with the elevated APOBEC-driven substitution patterns reported during sustained human-to-human transmission in recent mpox outbreaks.^4^

In contrast to clade IIa strains, clade IIb strains from Guinea clustered closely with sequences from the contemporaneous outbreak in Sierra Leone, indicating epidemiological links across national borders. In neighboring Sierra Leone, increasing numbers of mpox cases were reported starting in January 2025. The corresponding genomes were previously described as lineage G.1, a sub-lineage of A.2.^23^ In addition, the high number of APOBEC-driven substitutions in the clade IIb genomes suggests they originate from human-to-human transmission chains.

Our findings also emphasise the importance of combining decentralised diagnostic infrastructure with in-country sequencing capacity. Without diagnostic testing outside the capital region, samples representing distinct transmission dynamics would likely not have been identified, and without national sequencing capacity, the concurrent circulation of two epidemiologically distinct viral populations would have remained unrecognised. Together, this integrated approach has direct implications for public health decision-making by enabling context-specific responses that distinguish between local zoonotic emergence and sustained human-to-human transmission.

A limitation of our study is the uneven geographic sampling and the fact that genomes were obtained for only a subset of confirmed cases during the 2025 outbreak. Nevertheless, clear spatial and genomic structuring was observed, with distinct viral populations consistently associated with different diagnostic regions. Given that mpox cases were only confirmed in regions with functional laboratory infrastructure, it is tempting to speculate that more and more cases will be detected as laboratory and surveillance capacities expand geographically.

## Conclusion

In conclusion, we report mpox cases in Guinea arising from both independent zoonotic introductions (clade IIa) and sustained human-to-human transmission (clade IIb). Viral sequence data facilitate an effective use of public health capacities, as zoonotic spillover and sustained human transmission require different surveillance and control measures. The detection of cases only around the laboratory in Guéckédou suggests that a large number of cases still remain undetected and underscores the need to expand decentralized diagnostic capacity across larger geographic areas. In-country whole-genome sequencing capacity is important to inform the response to emerging viral threats.^27^

## Contributions

Conceived and designed the study: J.C., N.P.P., F.R.K., G.A., S.D., S.B.

Collected data and/or performed laboratory diagnostics: J.C., N.P.P., F.R.K., G.A., Kab.K., B.S., S.R., S.L.M., M.C., Kar.K., T.E.M., F.M.T., M.K., M.B.K.

Medical examinations and/or field investigations: S.A.B., F.T., O.T.

Performed sequencing and/or sequence validation: J.C., N.P.P., S.R., M.C.

Formal phylogenetic analysis: J.C., N.P.P., S.R., M.L., P.L.

Project implementation: J.C., N.P.P., F.R.K., G.A., S.R., E.V.N., C.v.G., E.K., S.C., A.C., S.G., S.D., S.B.

Funding acquisition: S.G., S.D., S.B.

Wrote the manuscript: J.C., N.P.P., F.R.K., G.A., S.D., S.B.

Edited the manuscript: all authors.

All authors read and approved the contents of the manuscript.

Edited the manuscript: all authors.

## Declaration of generative AI and AI-assisted technologies in the coding and writing process

ChatGPT was used to assist the coding process. All generated code was subsequently reviewed and validated by the developers to ensure accuracy and reliability. During the preparation of this work, the authors used ChatGPT in order to edit sentences. The authors reviewed and edited the content as needed and take full responsibility for the content of the publication.

