## Supplementary Appendix for "Emergence of mpox in Guinea: sporadic clade IIa cases and a clade IIb upsurge"

[a] Equal contribution first authors

[b] Equal contribution last authors

**Affiliations:**

### Tables

#### *Table S1*

**Mpox confirmed cases diagnosed at LFHV-GKD, Guéckédou (n=7), between 26 August 2024 and 31 December 2025, and their corresponding sequencing results.**

| Nr | ID | Age, y/sex | Prefecture | Date-sampling | Coverage [%\| | Clade | Genebank ID |
|---|---|---|---|---|---|---|---|
| 1 | G1027 | 6-10/F | Macenta | 30-Aug-2024 | 100 | IIa | PV993193 |
| 2 | G1041 | 21-25/F | Macenta | 4-Dec-2024 | 100 | IIa | PV993194 |
| 3 | G1104 | 41-45/M | Guéckédou | 17-Aug-2025 | 99 | IIa | PX782070 |
| 4 | G1100 | 31-35/F | Guéckédou | 22-Aug-2025 | 99 | IIa | PX782069 |
| 5 | G1101 | 6-10/M | Guéckédou | 30-Aug-2025 | na | na | na |
| 6 | G1115 | 16-20/M | Guéckédou | 29-Sep-2025 | 100 | IIa | PX990728 |
| 7 | G1116 | 16-20/M | Guéckédou | 29-Sep-2025 | na | na | na |

*Table S2*

**Characteristics of mpox confirmed cases in Guinea 2025 outbreak diagnosed at CRV, Conakry (n=728) – from 3 June to 29 December 2025.**

| Characteristics | Number (%) |
|---|---|
| **Gender** | |
| Female | 284 (39·0) |
| Male | 444 (61·0) |
| **Age (years)** | |
| Median (IQR25-75), Min-Max | 24·5 (16-32), 1-75 |
| **Age groups*** | |
| <16 | 173 (24·3) |
| 16-23 | 168 (23·6) |
| 24-32 | 207 (29·1) |
| 33-52 | 147 (20·6) |
| >52 | 17 (2·4) |
| **Region** | |
| Conakry | 570 (78·3) |
| Kindia | 109 (15·0) |
| Boké | 33 (4·5) |
| Kankan | 6 (0·8) |
| Faranah | 5 (0·7) |
| Mamou | 2 (0·3) |
| Boffa | 2 (0·3) |
| Dubreka | 1 (0·1) |
| **Month**** | |
| June | 5 (0·7) |
| July | 339 (46·6) |
| August | 189 (26·0) |
| September | 148 (20·3) |
| October | 42 (5·8) |
| November | 4 (0·5) |
| December | 1 (0·1) |

IQR : Interquartile range

*Total number differs from total as missing data (n=712)

** Based on sampling date

### Figures

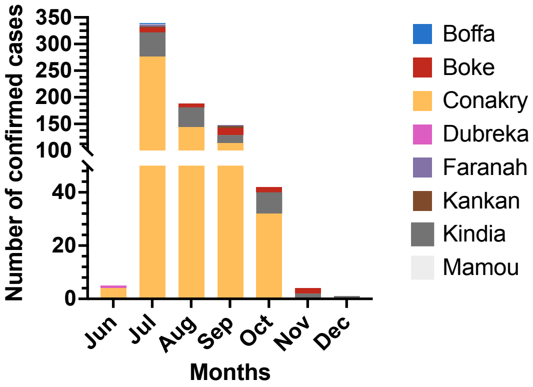

***Figure S1***

**Temporal distribution mpox cases diagnosed at CRV by region during the 2025 clade IIb outbreak in Guinea.**

Monthly distribution of laboratory-confirmed mpox cases diagnosed at CRV between 3 June and 29 December 2025 (n = 728), with bars stacked by region to illustrate temporal and regional trends in case number.
